# Beyond diversity: A whole system view of vegetation’s control of erosion

**DOI:** 10.64898/2026.10.08.757748

**Authors:** Janniry Cabrera Belen, Daniel N Livsey, Caroline Benfer-Martin, Tyler A Groh, Bradley J. Cardinale

**Author notes:** Corresponding author: Janniry Cabrera Belen.

## Abstract

Streambank erosion degrades both terrestrial and aquatic ecosystems, and riparian buffers are among the most widely used tools to control it. Yet most of what is known about how riparian vegetation controls erosion comes from one or two-species systems, which overlook the diversity of riparian communities and species interactions (competition and facilitation) that may shape the root systems that bind soil. We used a streambank mesocosm experiment to test whether plant species richness alters root traits, and whether root traits and species interactions reduce erosion. We established 85 mesocosms ranging from unvegetated controls to six-species polycultures, measured root biomass, length, volume, and diameter, quantified species interactions with relative yield totals (RYT), and measured erosion rate with a jet erosion test. Vegetated mesocosms eroded significantly less than unvegetated controls (*t* = 4.05, *p* = 0.023). Richness did not affect most root traits, although mean root diameter decreased as richness increased (*β* = -0.0225, *p* = 0.047). RYT ranged from competition (RYT < 1) to facilitation (RYT >1), yet neither interactions nor any root trait was directly associated with erosion rate. However, RYT was negatively associated with soil moisture (*β* = -0.196, *p* = 0.010). Piecewise structural equation models showed that erosion was best explained by a pathway linking species presence, root length, and soil moisture (AIC = 39.27; Fisher’s C = 11.27, *df* = 12, *p* = 0.506). *Panicum virgatum* increased root length, whereas *Lolium multiflorum* and *Echinochloa crus-galli* reduced it; longer roots were associated with drier soil (standardized *β* = -0.31, *p* = 0.005), and wetter soils eroded more slowly (standardized *β* = -0.39, *p* < 0.001). Which species were present mattered more than how many, and plants were linked to erosion through soil moisture rather than root traits alone. Species selection for riparian buffers may need to consider effects on soil moisture as well as rooting extent, particularly as droughts followed by intense storms become more common under a changing climate.

## INTRODUCTION

Streambank erosion is a growing environmental challenge driven by deforestation of riparian vegetation, unsustainable agricultural practices, and urbanization that have pushed contemporary erosion rates above geologic baselines (Wilkinson & McElroy, 2007; Montgomery, 2007). Its consequences span two ecosystems at once: soils underpin terrestrial productivity, and their loss reduces both the quantity and quality of vegetation an ecosystem can support (Pimentel et al., 1995), while the mobilized soil particles load stream systems with excess sediment and nutrients (Bilotta & Brazier, 2008). Terrestrial and aquatic consequences of streambank erosion converge in riparian zones, where vegetation regulates the movement of soil and nutrients between systems, governing both soil retention on land and water quality in streams (Naiman & Décamps, 1997; Riis et al., 2020), causing riparian buffers to be a widely used tool for protecting soils from erosion and filtering runoff (Osborne & Kovacic, 1993). The effectiveness of riparian buffers in terms of streambank erosion has traditionally been understood through physical characteristics (e.g. stream order, morphology, soil types) and the role that plant structures play in resisting it. Aboveground, leaves and stems intercept rainfall and slow flow across the soil surface; belowground, roots bind soil particles (Gyssels et al., 2005), reinforce soil strength (De Baets et al., 2008), and increase the resistance of soils to detachment (De Baets et al., 2011). The current understanding of the role of riparian plant communities in streambank erosion overlooks diversity and species interactions present in natural riparian communities, both of which may further shape erosion (Allen et al., 2014).

Past work has studied specific root traits of monocultures or two-species assemblages (Pollen-Bankhead & Simon, 2010) in relation to streambank erosion. For example, root diameter has been studied to quantify how tensile strength binds soil (De Baets et al., 2008), and root distribution and strength for bank reinforcement (Abernethy & Rutherfurd, 2001). Useful as these single to two species studies are for understanding particular root traits, they do not capture the diversity of riparian communities. Plant diversity has the capacity to influence streambank erosion through shaping root systems, greater species richness for example can change root biomass, rooting depth, and belowground allocation (Spehn et al., 2005; Mommer et al., 2010; Mueller et al., 2013; Ravenek et al., 2014), although these effects are often driven by species composition rather than richness itself (Siebenkäs & Roscher, 2016). Much of this root variation arises from how neighboring species interact belowground, and these interactions are a central mechanism structuring plant communities (Wilson & Tilman, 1991; Bruno et al., 2003; HilleRisLambers et al., 2012). Competition for space and resources can drive plants to explore the soil more extensively (Casper & Jackson, 1997; Gersani et al., 2001; Von Felten & Schmid, 2008; Lepik et al., 2021), while facilitation, a widespread positive interaction among plants (Bertness & Callaway, 1994; Callaway & Walker, 1997), can promote belowground growth (Yu et al., 2021). If diversity and species interactions shape the root traits that bind soil, then plant communities may influence erosion not only through the variety of root traits present, but through the changes in belowground structure that diversity itself produces.

Here we test how plant diversity, root traits, and interactions influence erosion, using a large-scale streambank mesocosm experiment. We established 85 mesocosms spanning a gradient of species richness (unvegetated controls, monocultures, two species, four species, and six species assemblages) and subjected each to a jet erosion test (JET) to quantify erosion rate.

We measured root biomass, length, volume, and diameter, and quantified species interactions (competition and facilitation) using relative yield totals. We tested three hypotheses: First, we hypothesized that as plant species richness increased so would belowground root traits. Second, we hypothesized that greater amounts of root traits like biomass and volume would lead to less erosion because denser root systems would bind soil particles and increase resistance to detachment. Third, we hypothesized that species interactions, such as facilitation would lead to less erosion, because facilitation would drive plants to produce more roots and increase soil binding.

## METHODS

### Greenhouse experimental design

We conducted this experiment in Mendel’s greenhouse in Bellefonte, Pennsylvania, a 20,000-square-foot plant growth facility owned by The Pennsylvania State University. Streambank mesocosms were housed in one of Mendel’s 600-square-foot bays. Mesocosms were constructed using Sterilite^®^ latching containers measuring 60.3 cm in length, 40.6 cm in width, and 34.3 cm in height (Figure 1A). To allow drainage, 80 1-cm diameter holes were drilled into the bottom of each container. The bottoms were then lined with Vigoro matrix grid landscape fabric to reduce soil loss during watering while maintaining drainage. Locally sourced screened topsoil was added to each container, lightly compacted, and shaped into a flat soil surface and a bank with a 1.5:1 horizontal:vertical slope (Allen et al., 2016). Each mesocosm was directly seeded and maintained at a density of 18 plants, while controls were maintained without any plants to observe how the soil alone eroded after being transported and shaped in the mesocosms.

**Figure 1.**
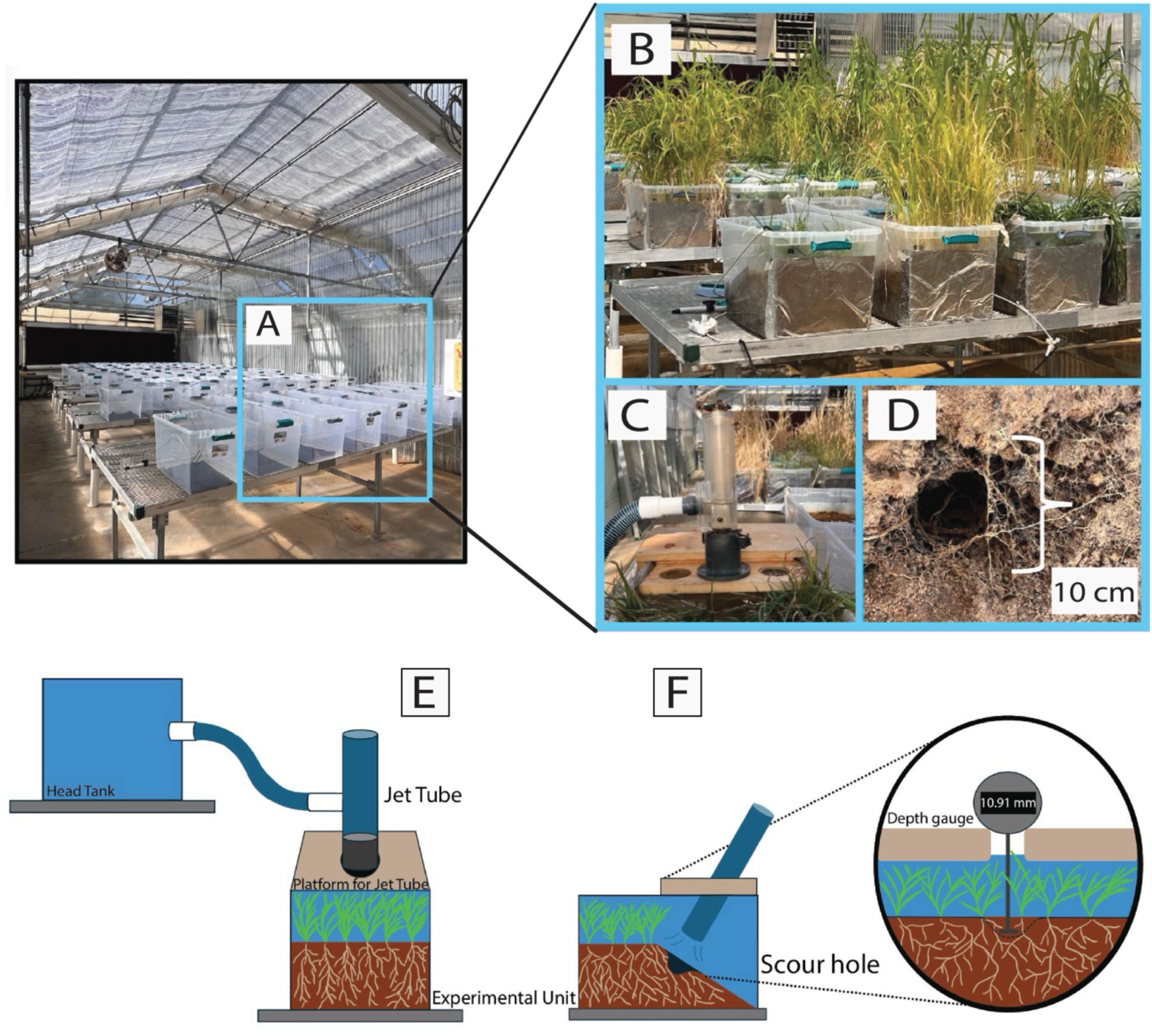
Diagram illustrating greenhouse experiment methods such as (A) set-up of experimental units, (B) growth of species in the experimental units, (C) the jet erosion test (JET) conducted in each unit, (D) the scour hole created from the JET that was done on the artificial banks slope to target root traits, (E) a front view of the JET set-up as the jet tube sits in the standardized platform and erodes the soil during the test, (F) a side view of the JET as the jet tube sits on the platform and erodes the slope to target root traits, as well as a view of the depth gauge meter that was utilized to measure the scour hole during the erosion test. Photographs by J. Cabrera Belen

During the growth period from July to October 2024, the slope was covered with Vigoro matrix grid landscape fabric and supported with a wooden slab to maintain the bank’s shape.

Species richness was manipulated using six native or naturalized herbaceous species common in Pennsylvania riparian areas (Rhoads & Block, 2007) and often used in stream restorations or as cover crops for erosion control (USDA-NRCS, 2000; Clark, 2007). Four C3 species were included: *Elymus virginicus* (native), *Agrostis gigantea* (naturalized), *Lolium multiflorum* (naturalized), and *Trifolium repens,‘Legacy’* (naturalized). Two C4 grasses were also included: *Panicum virgatum* (native) and *Echinochloa crus-galli* (naturalized). Species were directly seeded in July 2024 into the following treatments: controls (3 mesocosms), monocultures of each species (6 species × 3 replicates = 18 mesocosms), all possible biculture mixtures (15 bicultures × 3 replicates = 45 mesocosms), eight randomly selected four species mixtures (8 combinations × 2 replicates = 16 mesocosms), and one six species mixture (1 × 3 replicates = 3 mesocosms). For all treatments and replication, there were a total of 85 streambank mesocosms. Throughout the growth period, mesocosms were maintained by carefully removing weeds without disturbing the soil and were watered daily on the bank surface until saturated, with excess running off the bank and filtered out of the container. Greenhouse temperatures were maintained at 24 °C during the day and 18 °C at night. Artificial lighting was not used, but automatic shade cloths were deployed during midday to reduce excessive heat within the bay.

### Jet erosion test (JET)

After the three-month growth period in October 2024, a single submerged jet erosion test (JET) was conducted in each streambank mesocosm (Figure 1C). The jet erosion device (JED) is a standard tool used in erosion research and regulated by the American Society for Testing and Materials (ASTM). It consists of a jet tube with a standard nozzle opening of 6.4 mm that regulates static pressure and the force of the water stream to erode soil (ASTM, 1999; Hanson & Cook, 2004). The jet tube was connected to a head tank positioned at a constant height above the mesocosms, which provided consistent static pressure and generated force of 2 kPa across all trials, a common force exhibited in streams. The JET was performed on the bank’s slope to isolate the effects of root traits on erosion (Figure 1D). Prior to the test, a soil core was collected so that soil covariates such as moisture, bulk density, and porosity could be included in subsequent analyses. Mesocosms were then flooded, and a standardized platform was placed over each container (Figure 1E). This platform ensured a consistent jet tube height above the soil surface and allowed a depth gauge to measure the distance between the platform and the soil surface before and during the test (Figure 1F). Each JET lasted 20 minutes, at the start of the test the JED was inserted into the platform and ran for predetermined time intervals. At each measurement point, the JED was removed, which paused the test and stopped erosion, and a depth gauge was placed on the platform to measure the scour hole (Figure 1F). These measurements were used to calculate erosion rates (mm per minute). Only one erosion test was possible per experimental unit due to container size, implications of this are highlighted in the discussion.

### Plant harvest and root scanning

Following the erosion tests, plants were harvested. Plant samples were stored at -20 °C until they could be cleaned and processed. Before scanning and measurement, roots were thoroughly cleaned of soil and debris. Roots were then arranged for scanning and scanned with an Epson Perfection 4490 flatbed scanner at 1200 dpi. Images were analyzed in RhizoVision Explorer v2.0.3 (Seethepalli & York, 2020) using algorithms described by Seethepalli et al. (2021). The software can quantify a multitude of root traits, including length, volume, and diameter. After scanning, samples were dried at 65 °C for 24 h. Roots and aboveground tissues were then weighed to determine biomass.

### Relative Yield of Roots

To quantify how interactions among plant species impact belowground root growth, we calculated relative yield totals (RYT). RYT is a standard metric used to infer whether species growing together in a mixed polyculture experience competition (RYT < 1) or facilitation (RYT > 1) (Loreau, 1998). Relative yield for each species was calculated by dividing its root biomass in polyculture by its monoculture root biomass, and species-level relative yields were then summed to obtain experimental unit-level RYT. Because monoculture root biomass varied substantially among individual plants, we estimated these metrics using an individual-based bootstrap procedure in R statistical software version 4.6.1 (R Core Team 2026). For each species, the monoculture reference pool consisted of 54 plants drawn from three monoculture experimental units, and during each bootstrap iteration, we randomly sampled 18 plants with replacement from that pool to represent the biomass of one monoculture experimental unit. This resampling procedure was repeated 1,000 times, and the resulting distributions were used to calculate the mean and standard deviation of species-level relative yields and experimental unit-level RYT values.

### Statistical analyses

To test our hypotheses, we analyzed the effects of plant species richness on root traits (biomass, volume, length, and diameter) and erosion rate, as well as the effects of root traits and interactions (competition and facilitation) on erosion rate utilizing generalized linear mixed models (GLMMs) fit with the package *glmmTMB* (Brooks et al., 2017; McGillycuddy et al., 2025) in R. We also conducted a Welch’s two-sample t-test to compare the erosion rates between vegetated mesocosms and unvegetated controls. Before analysis, all variables were checked for normality, and right-skewed root traits (biomass, volume, and length) were natural-log transformed. We used gaussian GLMMs with an identity link to test the effects of plant species richness on each root trait (biomass, volume, length, and mean diameter) and on erosion rate, and the effects of root biomass, root volume, and RYT on erosion rate. RYT models included only polycultures (n = 64) and treated RYT as a continuous predictor. Because soil moisture content is a proportion bounded between 0 and 1, we tested the effect of RYT on soil moisture with a beta GLMM and a logit link. Mesocosms were arranged across five greenhouse tables that shared light and temperature conditions, so all GLMMs included table as a random intercept to account for variation caused by greenhouse table placement. For all models, visual inspection of residuals occurred with the package *DHARMa* (Hartig, 2026), if quantile deviations were detected normality was double checked utilizing a Shapiro-Wilk test.

Since erosion is likely governed by interacting biological and soil processes rather than single predictors, we used piecewise structural equation models (pSEMs) fit with the package *piecewiseSEM* (Lefcheck, 2016). Unlike traditional SEM, which estimates all relationships simultaneously from a single covariance matrix, pSEM breaks down pathways into a set of component models. Our pSEMs structure was based on established mechanisms and exploratory analysis linking species to roots, soil, and erosion. Each pSEM was comprised of three component models: (1) how species presence affects the root trait, (2) the relationship between the root trait and soil moisture content, and (3) how the focal root trait and soil moisture content affects erosion rate. The component models were fit using linear models (*lm*) from the R base *stats* package and linear mixed-effects models (*lmer*) from the *lme4* package (Bates et al., 2015). Each root pSEM included the following candidate variables: the presence of all plant species (*Elymus virginicus*, *Agrostis gigantea*, *Lolium multiflorum*, *Trifolium repens* ‘Legacy’, *Panicum virgatum*, and *Echinochloa crus-galli*), the root trait of interest (biomass, length, volume, or diameter), soil properties (bulk density, porosity, and moisture content), and erosion rate. Non-significant predictors (*p* > 0.05) were removed sequentially, beginning with the highest *p*-value, until AIC decreased by less than 2 points. We used tests of directed separation to identify missing paths and added them where biologically plausible, which added direct species to soil moisture paths to the root volume and diameter models. We compared the four pSEMs using AIC derived from Fisher’s C (Shipley, 2013; Lefcheck, 2016) and considered the model with the lowest AIC the best supported. Global fit was evaluated with Fisher’s C, for which a non-significant *p*-value (*p* > 0.05) indicates no evidence of missing paths. We report standardized path coefficients extracted with the *coefs* function and calculated indirect effects as the product of the standardized coefficients along each pathway.

## RESULTS

### Richness, root traits, and soil effects on erosion

A *t*-test showed that vegetated mesocosms had significantly lower erosion rate than unvegetated mesocosms (*t* = 4.05, *df* = 3.25, *p* = 0.023). In this experiment, plant species richness did not cause differences in most root traits (*p* > 0.05), the only root trait to be statistically different was mean root diameter (*β* = -0.0225, SE = 0.0114, *z* = -1.983, *p* = 0.047) (Figure 2D). Plant species richness also did not cause differences in erosion rate (Figure 2E) (*p* > 0.05). Neither root biomass nor volume was associated with erosion rate (Figure 3 A & B). Among polycultures, RYT was not associated with erosion rate (Figure 3C). See Appendix S1: Tables S1 and S2 for full model results.

**Figure 2.**
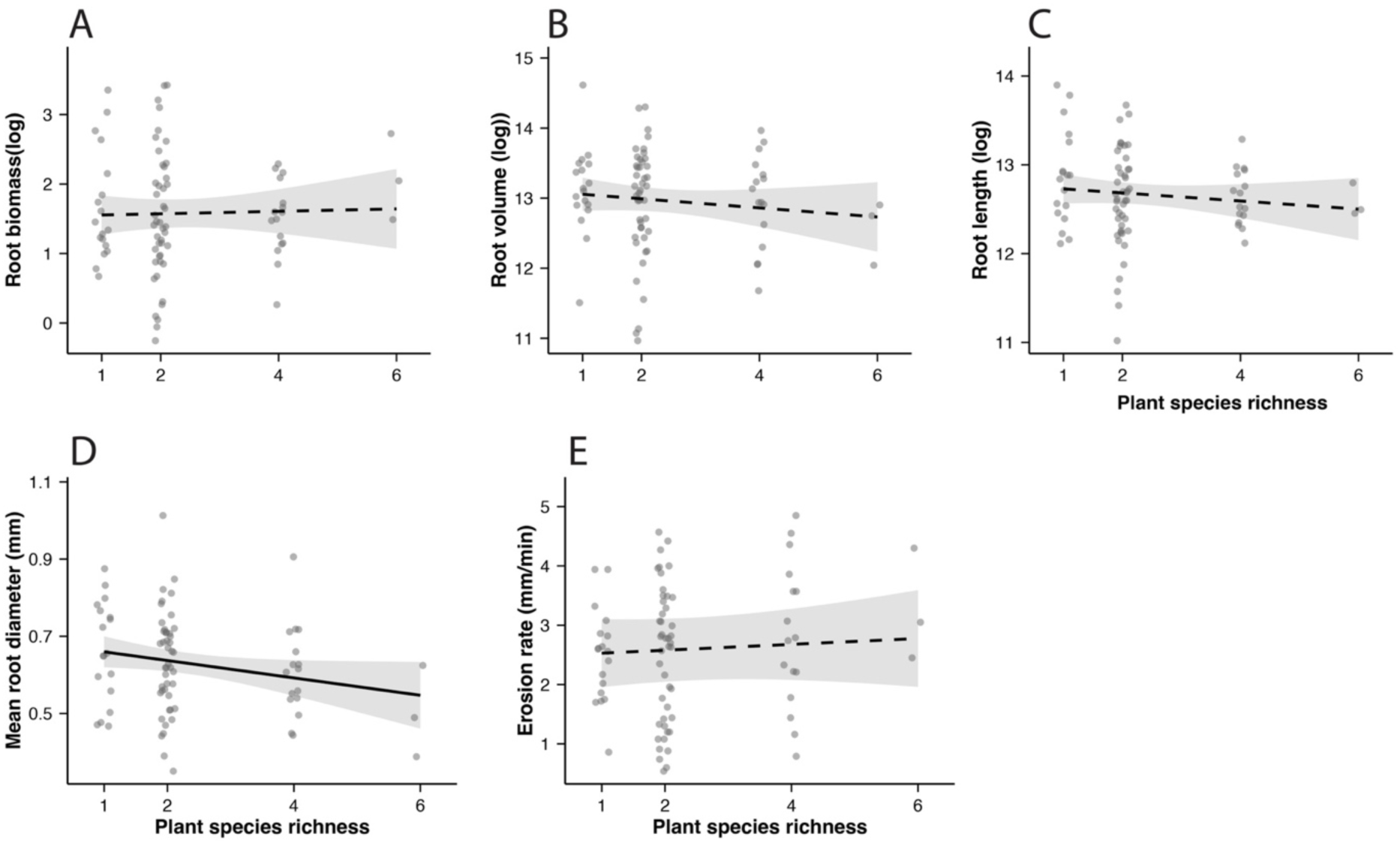
Effects of plant species richness on root traits and erosion rate: (A) root biomass (g), (B) root volume (mm³), (C) root length (mm), (D) mean root diameter (mm), and (E) erosion rate (mm/min). Root biomass, volume, and length were natural-log transformed. Data points represent each mesocosm (n = 82). Lines are derived from the GLMM with shaded 95% confidence intervals. The solid line indicates a significant effect of species richness (p < 0.05), and dashed lines indicate non-significant effects.

**Figure 3.**
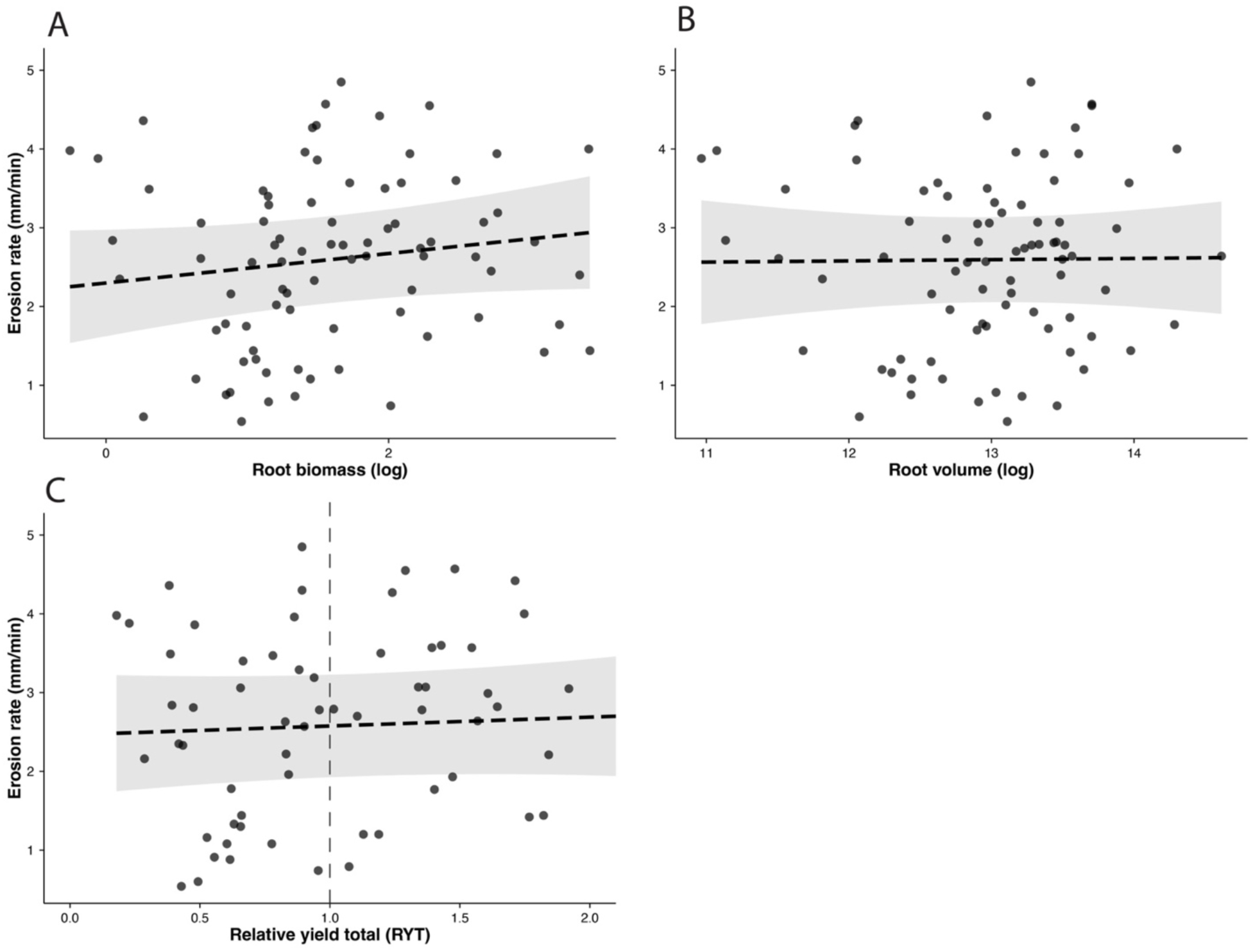
Effects of root traits and species interactions on erosion rate: (A) root biomass (g), (B) root volume (mm³), and (C) relative yield total (RYT) of polyculture mesocosms. Root biomass and volume were natural-log transformed. Data points represent each mesocosm (A, B: n = 82; C: n = 64 for polycultures). Lines are derived from the GLMMs with a shaded 95% confidence intervals; dashed lines indicate non-significant effects (p > 0.05). In (C), the dotted vertical line marks RYT = 1 the expected yield for monocultures; values below 1 indicate competition and values above 1 indicate facilitation.

### Piecewise structural equation models (pSEM)

During exploratory analysis, a pattern between root traits such as root length and moisture content in the context of erosion rate became apparent (Figure 4A), greater amounts of root length were correlated with soil moisture, and soil moisture was associated with erosion rates. Additionally, RYT was negatively associated with soil moisture content (*β* = -0.196 on the logit scale, SE = 0.076, *z* = -2.582, *p* = 0.010; Figure 4B). All four models had good fit with the data (Fisher’s C, p = 0.24-0.74; Table 1) and had no evidence of missing paths. Of the four pSEMs, the root length model was best supported (AIC = 39.27; Table 1). In the root length model (Fisher’s C = 11.27, *df* = 12, *p* = 0.506; Figure 5), species presence explained 33% of the variation in root length (marginal *R²* = 0.33, conditional *R²* = 0.38). *Panicum virgatum* increased root length (*β* = 0.40, SE = 0.09, *t* = 4.20, *p* < 0.001), whereas *Echinochloa crus-galli* (*β* = -0.25, SE = 0.09, *t* = -2.64, *p* = 0.010) and *Lolium multiflorum* (*β* = -0.41, SE = 0.10, *t* = -4.28, *p* < 0.001) reduced it. Root length was negatively associated with soil moisture content (*β* = -0.023, SE = 0.008, *t* = -2.92, *p* = 0.005; standardized *β* = -0.31; *R²* = 0.10). Soil moisture content was negatively associated with erosion rate (*β* = -11.09, SE = 2.51, *t* = -4.42, *p* < 0.001; standardized *β* = -0.39), whereas root length had no direct association with erosion (*β* = -0.065, SE = 0.19, *t* = -0.34, *p* = 0.738; marginal *R²* = 0.13, conditional *R²* = 0.50). Indirect effects of species on erosion through root length and soil moisture were small: +0.05 for *P. virgatum*, -0.05 for *L. multiflorum,* and -0.03 for *E. crus-galli* (Appendix S1: Tables S3-S6, Figures S1-S3).

**Figure 4.**
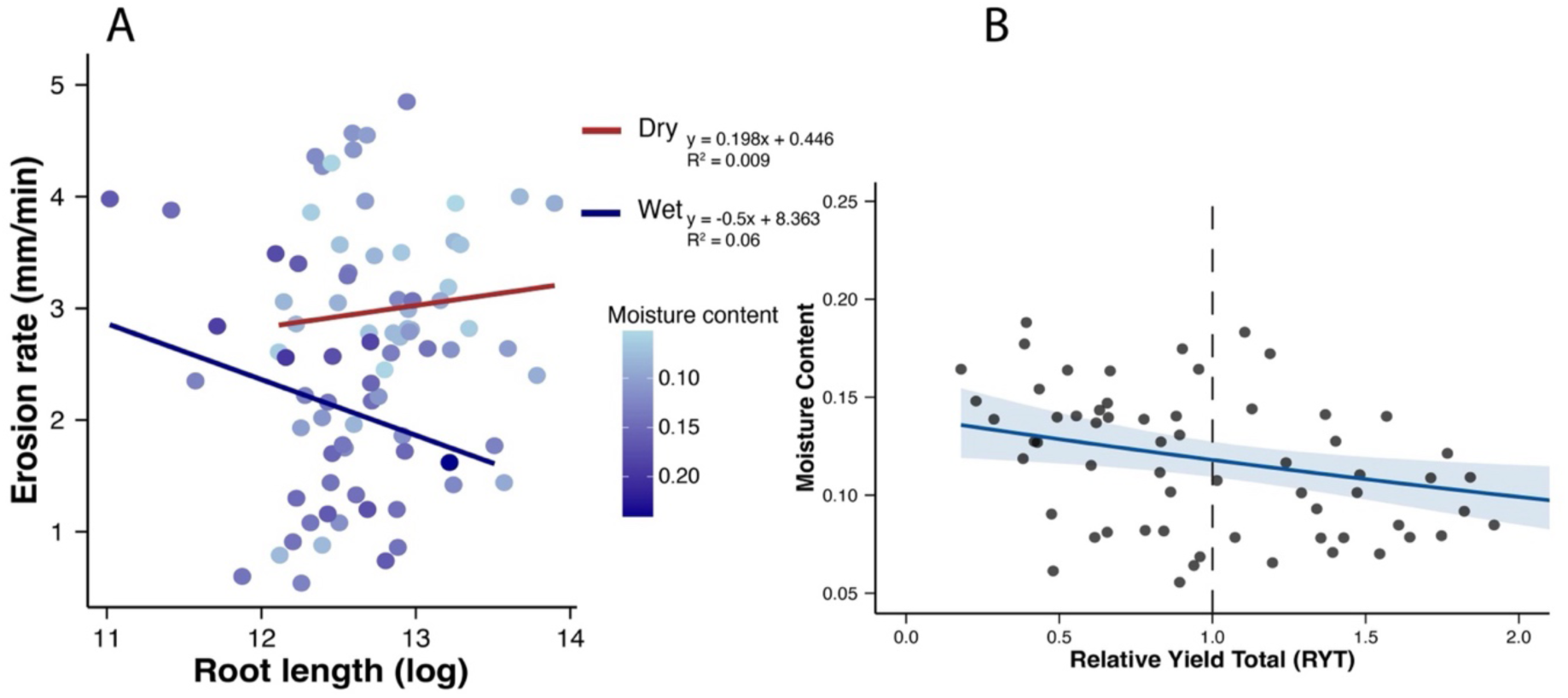
Relationships among root length, species interactions, soil moisture content, and erosion rate: (A) root length (mm; natural-log transformed) and erosion rate (mm/min), with data points colored by soil moisture content and separate lines for mesocosms classified as “wet” or “dry”. (B) Soil moisture content and relative yield total (RYT) of polyculture mesocosms (n = 64). The solid line is the fitted GLMM with shaded 95% confidence interval, indicating a significant negative effect of RYT on soil moisture content (p < 0.05). The dotted vertical line marks RYT = 1 the expected yield from monocultures; values below 1 indicate competition and values above 1 indicate facilitation.

**Figure 5.**
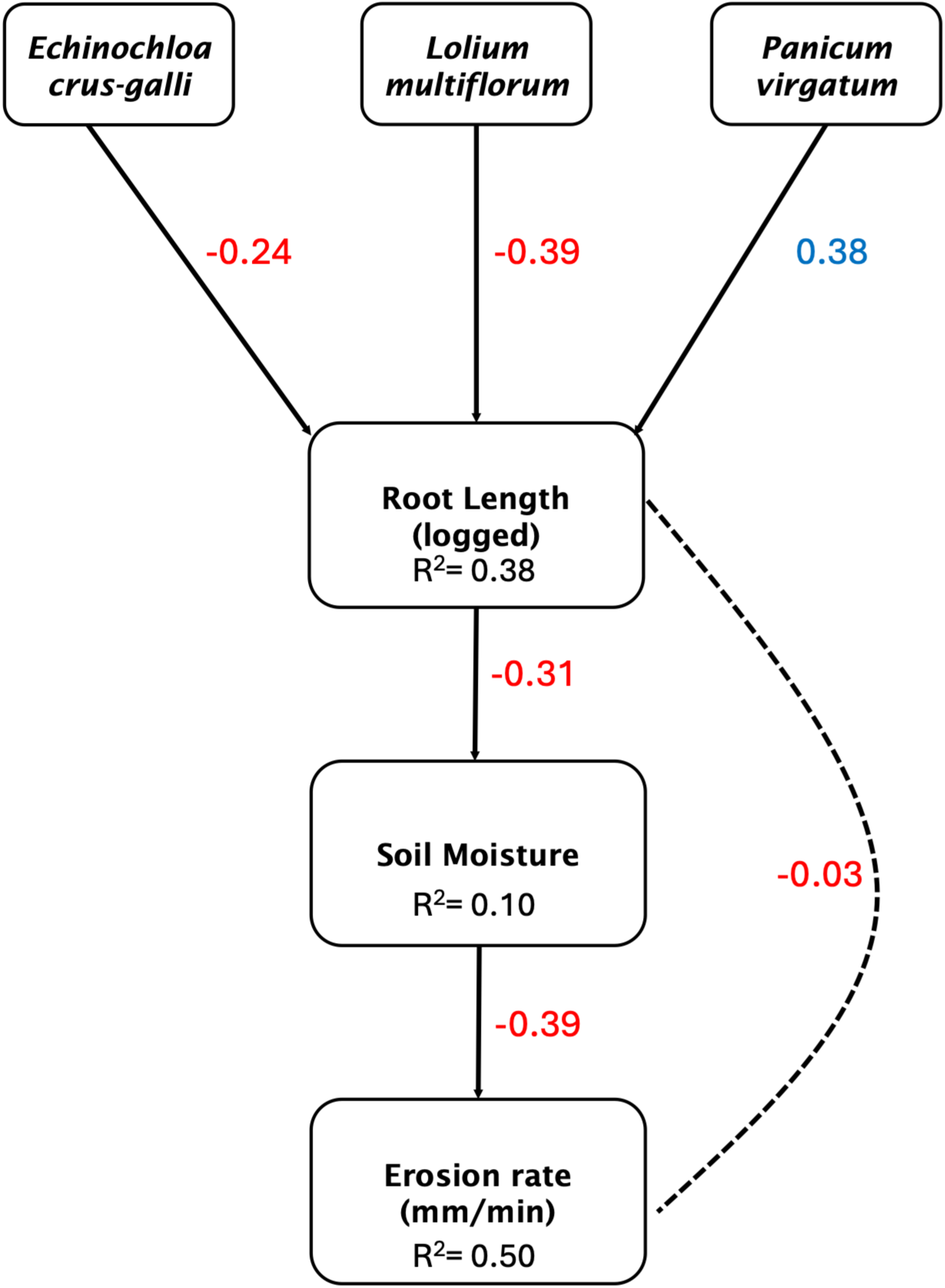
Piecewise structural equation model (pSEM) relating species presence, root length, soil moisture content, and erosion rate (mm/min) across all mesocosms (n = 82). Of the four candidate pSEMs, one for each root trait (Table 1), the root length model was selected because it had the lowest AIC and was the only one with a significant pathway from the root trait to soil moisture. Arrows point from predictor to response, and the numbers on the arrows are standardized path coefficients. Solid arrows are significant paths (p < 0.05, from the component lms or lmers), and the dashed arrow is a non-significant path. Positive standardized coefficients are shown in blue and negative paths in red. The presence of *Panicum virgatum* was associated with greater root length, while the presence of *Echinochloa crus-galli* and *Lolium multiflorum* was associated with lower root length. Greater root length was associated with lower soil moisture, and higher soil moisture was associated with lower erosion rates. R² values are shown for each response variable and model fit was assessed with Fisher’s C (11.27) and its *p* value (0.506), indicating that the model is a good fit of the data with no evidence of missing paths.

**Table 1.**
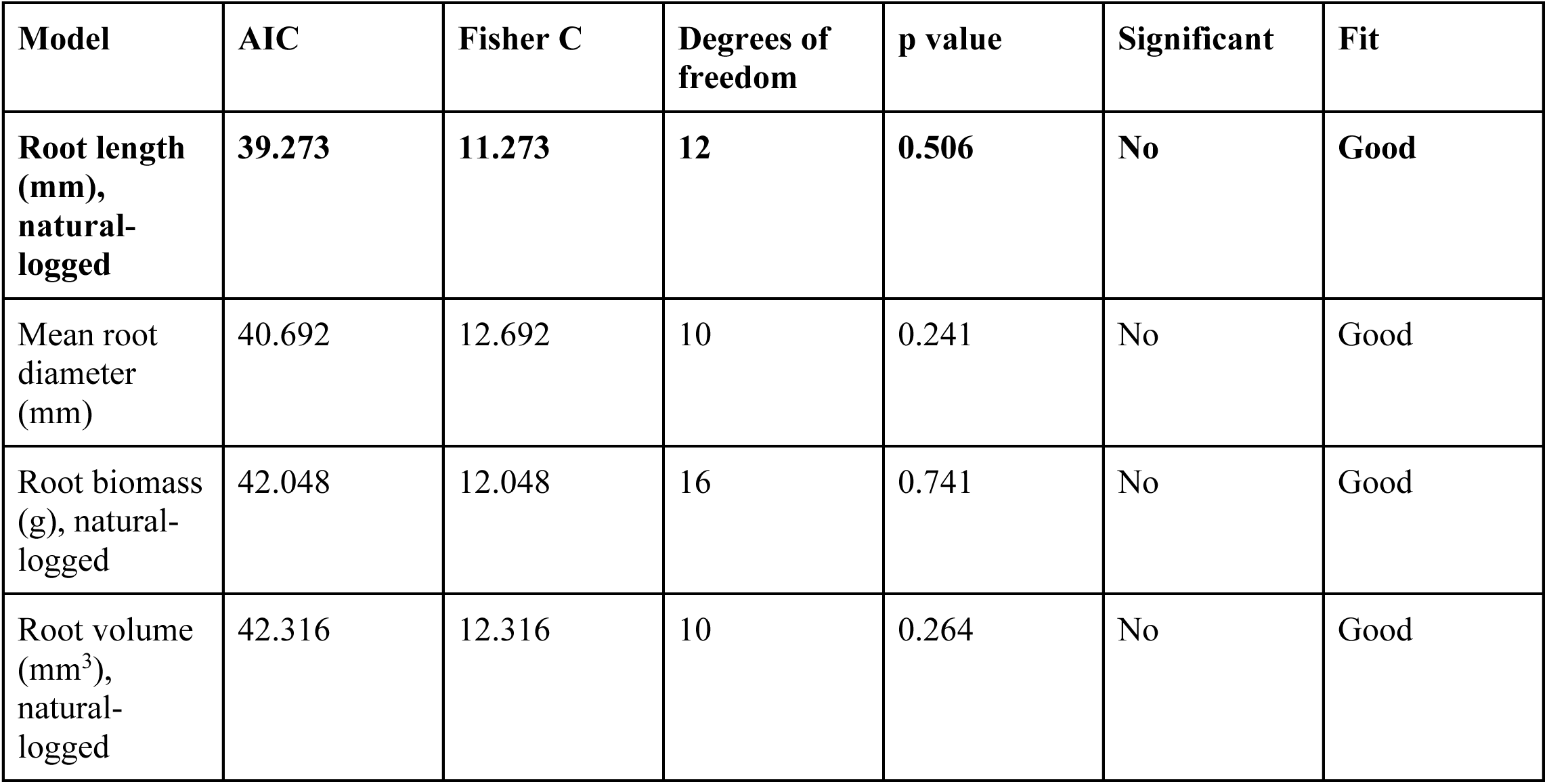
Comparison of piecewise structural equation models (pSEMs) for each root trait. Each model tested how species presence affects erosion rate through that root trait and soil moisture content (n = 82 mesocosms). Models are ranked by AIC calculated from Fisher’s C. Fisher’s C tests model fit, and a non-significant p-value (p > 0.05) indicates that the model is a good fit of the data, with no missing paths.

| Model | AIC | Fisher C | Degrees of freedom | p value | Significant | Fit |
| --- | --- | --- | --- | --- | --- | --- |
| **Root length (mm), natural-logged** | **39.273** | **11.273** | **12** | **0.506** | **No** | **Good** |
| Mean root diameter (mm) | 40.692 | 12.692 | 10 | 0.241 | No | Good |
| Root biomass (g), natural-logged | 42.048 | 12.048 | 16 | 0.741 | No | Good |
| Root volume (mm<sup>3</sup>), natural-logged | 42.316 | 12.316 | 10 | 0.264 | No | Good |

## DISCUSSION

In this experiment, we investigated how plant species richness shapes belowground root traits, and whether greater root biomass, root volume, and species interactions would reduce streambank erosion. Richness significantly affected only one root trait, mean root diameter, which decreased as richness increased. Neither root traits nor species interactions (RYT) were directly associated with erosion rate. However, RYT was negatively associated with soil moisture, and soil moisture was the strongest predictor of erosion. Piecewise structural equation models showed that erosion in our experiment is best understood as a product of the whole mesocosm system: the presence of particular species shaped root length, root length was associated with soil moisture, and soil moisture strongly determined how much soil eroded.

Although we did not detect richness effects on most root traits, the range of RYT values indicates that species interacted belowground in polycultures. Facilitative interactions (RYT >1) were associated with lower soil moisture indicating that species interactions may therefore influence erosion indirectly, through their effects on soil (Figure 4B). Additionally, root diameter declined with increasing richness (Figure 2D), whereas root characteristics in other experiments have been driven by functional composition rather than species richness (Siebenkäs & Roscher, 2016), suggesting that this response may depend on the particular species in our mixtures. The lack of response in our other root traits is consistent with biodiversity studies of root production, which have found mixed results, including no effect of richness on root production in some cases (Schmid et al., 2009) and effects that emerge only over time (Ravenek et al., 2014). Although we did not detect a direct effect of root biomass or volume on erosion (Figure 3A & B), vegetated mesocosms eroded less than unvegetated controls, consistent with the established protective role of roots (Gyssels et al., 2005; De Baets et al., 2006; Vannoppen et al., 2015). We attribute the absence of root and interaction effects to the sampling scale of the jet erosion test. Our measures of root traits and interactions (RYT) integrate the root systems of the entire mesocosm, whereas each jet erosion test measured erosion rate at a single location on the bank surface. Because root distributions in soil are spatially heterogeneous (Abernethy & Rutherfurd, 2001; Wynn et al., 2004), the roots beneath the jet may not have been representative of the mesocosm root system. We recommend that future studies use multiple jet erosion tests per experimental unit to better capture the spatial heterogeneity of root systems when evaluating how plant communities influence erosion.

Of our four piecewise structural equation models, the root length model was the best supported (Figure 5, Table 1). The root diameter model was within 2 AIC units, but we present the root length model because it had a significant pathway from the root trait to soil moisture. In this model, erosion was shaped by the species rather than richness. *Panicum virgatum* increased root length, whereas *Lolium multiflorum* and *Echinochloa crus-galli* reduced it. Longer roots were associated with lower soil moisture, and wetter soils eroded substantially less, consistent with the patterns of soil moisture and erosion in the literature (Livsey et al., 2025). *Panicum virgatum* had a small positive indirect effect on erosion (standardized coefficients: 0.38 × -0.31 × -0.39 ≈ +0.05), its longer roots were associated with drier soil and faster erosion. *Echinochloa crus-galli* (≈ -0.03) and *Lolium multiflorum* (≈ -0.05) had negative indirect effects. While these effects are small, they indicate that the species producing the most extensive root systems did not produce the most erosion-resistant banks in our experiment. Species with longer roots may instead contribute more to bank strength in erosion processes like mass failure than to resistance against fluvial erosion, the process we measured. The direction of the root-moisture relationship also cannot be resolved with our design, it is possible that longer roots may have decreased soil moisture (Cai et al., 2022), but naturally drier mesocosms could equally have stimulated root elongation as plants foraged for water (Comas et al., 2013). We therefore interpret this pathway as an association that identifies soil moisture as the strongest predictor variable of erosion in our system, with species identity influencing both root length and soil moisture.

Our results add to a small but growing body of work linking biodiversity to erosion; our work is amongst the first to consider plant community interactions as an influencing factor of erosion. Previous work with macroinvertebrates has shown that interactions among caddisfly species influence sediment binding (Albertson et al., 2014), and our results suggest that plant species interactions may influence erosion indirectly, through their association with soil moisture. The most direct comparison to our study is Allen et al. (2016), who used a jet erosion test in streambank mesocosms and found that plant species richness reduced erosion, but only at low richness levels, with polycultures not outperforming the best monocultures. Similarly, Berendse et al. (2015) found that net soil loss doubled when plant diversity declined, largely because diverse communities compensated when individual species were lost under environmental stress. Additionally, diverse plant communities also improve soil physical properties through complementary root traits (Gould et al., 2016), and in the Jena Experiment, the presence of particular plant functional groups altered soil water content in addition to species richness (Fischer et al., 2019). Our results connect these lines of evidence by suggesting that community composition can affect erosion through its effect on soil moisture, a pathway that biodiversity-erosion studies have rarely considered alongside root traits.

Species selection for riparian buffers typically emphasizes rapid establishment or general conservation seed mixes from commercial vendors (e.g., Ernst Conservation Seeds), with the expectation that extensive root systems protect banks from erosion. Our results suggest that, by drying the soil, these species may instead increase fluvial erosion. But they may be advantageous where geotechnical processes such as mass failure dominate, because they can increase matric suction and stabilize banks (Simon & Collison, 2002; Pollen-Bankhead & Simon, 2010). Practitioners may therefore need to weigh a species’ effect on soil moisture alongside its rooting extent, particularly where fluvial erosion rather than mass failure is the dominant process. This trade-off between roots and moisture will become more pressing as precipitation regimes shift toward more frequent intense storms and longer droughts (Knapp et al., 2008). Furthermore, root responses to drought can shift roots to deeper soil layers (Comas et al., 2013), or, under prolonged drought, reduce root length in grasses (Sun et al., 2024). Shifts in roots may further alter their ability to protect banks against both fluvial and geotechnical erosion. Because root length and soil moisture were linked in our system, and moisture was the strongest predictor of erosion, drought-driven changes in root systems may alter streambank erosion in ways that are difficult to predict. Understanding how plant species, diversity, and interactions shape both root traits and soil moisture will be essential for designing riparian buffers that remain effective in an increasingly variable climate.

## ACKNOWLEDGMENTS

We thank Sam Gruneberg and Mendel’s Greenhouse staff for greenhouse space and research support. We thank Maya Fulton and Sae Clausing for help with experimental maintenance, as well as the Stillwater Oklahoma USDA-ARS Unit for guidance on jet erosion tests. We thank Danilo Ferreira Borges dos Santos and Dr. Shannon Buttimer for help with statistical models and feedback on statistical analyses. We thank Dr. Bradley J. Cardinale for funding this work.

## AUTHOR CONTRIBUTIONS

JCB and BJC conceived and designed the experiment, BJC funded the experiment, DNL trained and advised JCB on designing the erosion test portions of the experiment. JCB and CBM collected the data; DNL advised on erosion test data collection. JCB analyzed and visualized data with help from TAG, DNL, and BJC. JCB wrote the first draft; all authors contributed to editing the article.

