## Appendix S1 for "Beyond diversity: A whole system view of vegetation’s control of erosion"

**Table S1:** Generalized linear mixed models (GLMMs) of plant species richness effects on root traits and erosion rate.

| Predictor | Response | Estimate | Std. Error | z value | p value |
|---|---|---|---|---|---|
| Plant species richness | Root biomass (g), natural-log transformed | 0.0176 | 0.0744 | 0.237 | 0.813 |
| Plant species richness | Root length (mm), natural-log transformed | -0.0456 | 0.0458 | -1.0 | 0.319 |
| Plant species richness | Root volume ($mm^3$), natural-log transformed | -0.0648 | 0.0647 | -1.0 | 0.317 |
| Plant species richness | Mean root diameter (mm) | -0.0225 | 0.0114 | -1.983 | **0.047** |
| Plant species richness | Erosion rate (mm/min) | 0.0494 | 0.0863 | 0.573 | 0.567 |

All models: Gaussian GLMMs with greenhouse table as a random intercept (n = 82 planted mesocosms).

**Table S2:** Generalized linear mixed models (GLMMs) of root and interaction effects on erosion rate.

| Predictor | Response | Estimate | Std. Error | z value | p value |
|---|---|---|---|---|---|
| Root biomass (g), ln-transformed | Erosion rate (mm/min) | 0.1932 | 0.1275 | 1.516 | 0.130 |
| Root volume ($mm^3$), ln-transformed | Erosion rate (mm/min) | 0.0155 | 0.1451 | 0.107 | 0.915 |
| Relative yield total (RYT) | Erosion rate (mm/min) | 0.1123 | 0.1942 | 0.578 | 0.563 |

Root trait models: n = 82 planted mesocosms; the root biomass model included a dispersion submodel with ln root biomass as a predictor. RYT model: polycultures only (n = 64), with RYT as a continuous predictor. All models: Gaussian GLMMs with greenhouse table as a random intercept.

**Table S3a:** Linear mixed effects model (LMER) of species presence effects on root length for the first component model for the piecewise structural equation model (pSEM).

| Predictor | Response | Estimate | Std. Error | DF | t value | p value |
|---|---|---|---|---|---|---|
| *Echinochloa crus-galli* | Root length (mm), natural-log transformed | -0.2500 | 0.0946 | 77.6 | -2.643 | **0.010** |
| *Lolium multiflorum* | Root length (mm), natural-log transformed | -0.4085 | 0.0955 | 77.5 | -4.278 | **< 0.001** |
| *Panicum virgatum* | Root length (mm), natural-log transformed | 0.3963 | 0.0944 | 75.6 | 4.197 | **< 0.001** |

**Table S3b:** Linear model (LM) of root length effect on soil moisture for the second component model for the piecewise structural equation model (pSEM).

| Predictor | Response | Estimate | Std. Error | DF | t value | p value |
|---|---|---|---|---|---|---|
| Root length (mm), ln-transformed | Soil moisture content | -0.0233 | 0.0080 | 80 | -2.919 | **0.005** |

**Table S3c:** Linear mixed effects model (LMER) of moisture content and root length effects on erosion rate for the third component model for the piecewise structural equation model (pSEM).

| Predictor | Response | Estimate | Std. Error | DF | t value | p value |
|---|---|---|---|---|---|---|
| Soil moisture content | Erosion rate (mm/min) | -11.09 | 2.51 | 75.2 | -4.421 | **< 0.001** |
| Root length (mm), ln-transformed | Erosion rate (mm/min) | -0.0647 | 0.1925 | 75.6 | -0.336 | 0.738 |

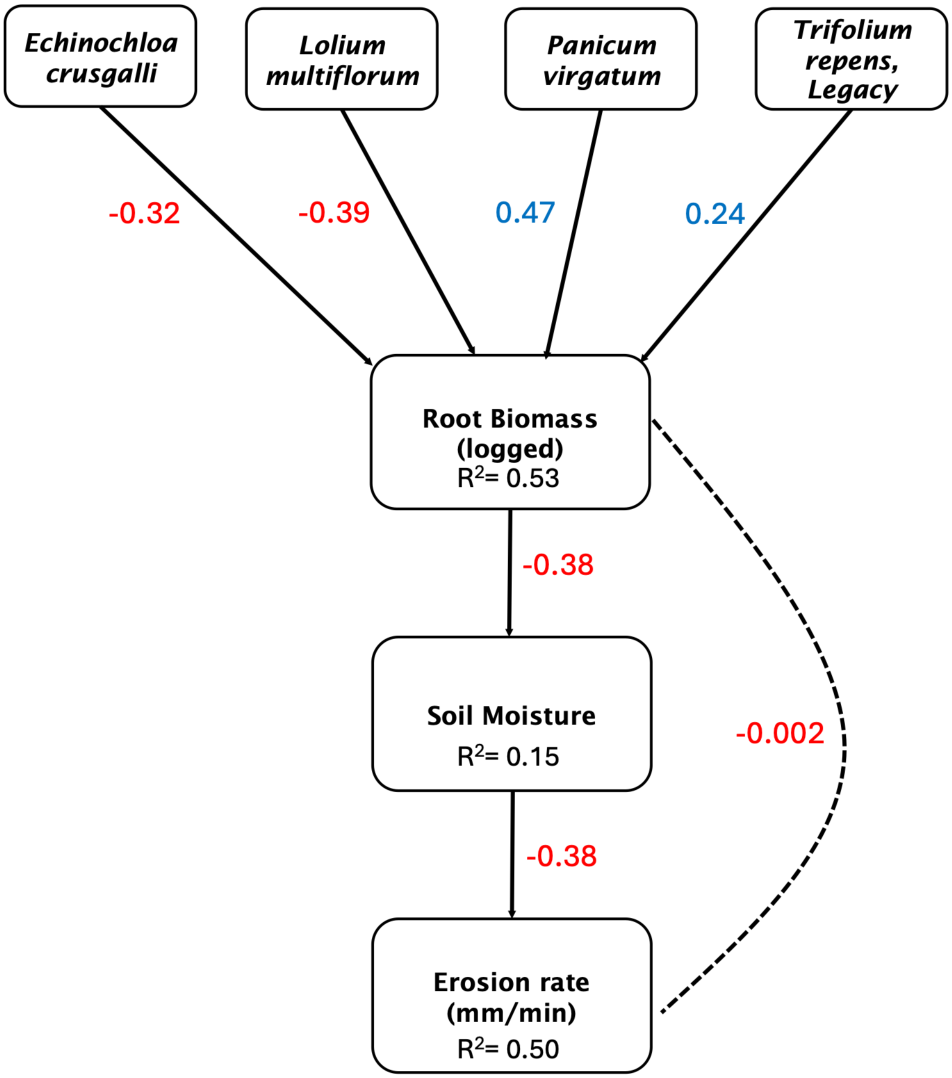


**Figure S1:** Visual for piecewise structural equation model (pSEM) for root biomass.

**Table S4a:** Linear mixed effects model (LMER) of species presence effects on root biomass for the first component model for the piecewise structural equation model (pSEM).

| **Predictor** | **Response** | **Estimate** | **Std. Error** | **DF** | **t value** | **p value** |
|---|---|---|---|---|---|---|
| *Echinochloa crus-galli* | Root biomass (g), natural-log transformed | -0.5371 | 0.1307 | 77.0 | -4.108 | **< 0.001** |
| *Lolium multiflorum* | Root biomass (g), natural-log transformed | -0.6476 | 0.1316 | 76.9 | -4.919 | **< 0.001** |
| *Panicum virgatum* | Root biomass (g), natural-log transformed | 0.8075 | 0.1306 | 75.1 | 6.184 | **< 0.001** |
| *Trifolium repens* 'Legacy' | Root biomass (g), natural-log transformed | 0.4188 | 0.1325 | 77.0 | 3.159 | **0.002** |

**Table S4b:** Linear model (LM) of root biomass effect on soil moisture for the second component model for the piecewise structural equation model (pSEM).

| **Predictor** | **Response** | **Estimate** | **Std. Error** | **DF** | **t value** | **p value** |
|---|---|---|---|---|---|---|
| Root biomass (g), ln-transformed | Soil moisture content | -0.0181 | 0.0048 | 80 | -3.780 | **< 0.001** |

**Table S4c:** Linear mixed effects model (LMER) of moisture content and root biomass effects on erosion rate for the third component model for the piecewise structural equation model (pSEM).

| **Predictor** | **Response** | **Estimate** | **Std. Error** | **DF** | **t value** | **p value** |
|---|---|---|---|---|---|---|
| Soil moisture content | Erosion rate (mm/min) | -10.86 | 2.58 | 75.1 | -4.206 | **< 0.001** |
| Root biomass (g), ln-transformed | Erosion rate (mm/min) | -0.0029 | 0.1236 | 75.7 | -0.023 | 0.981 |

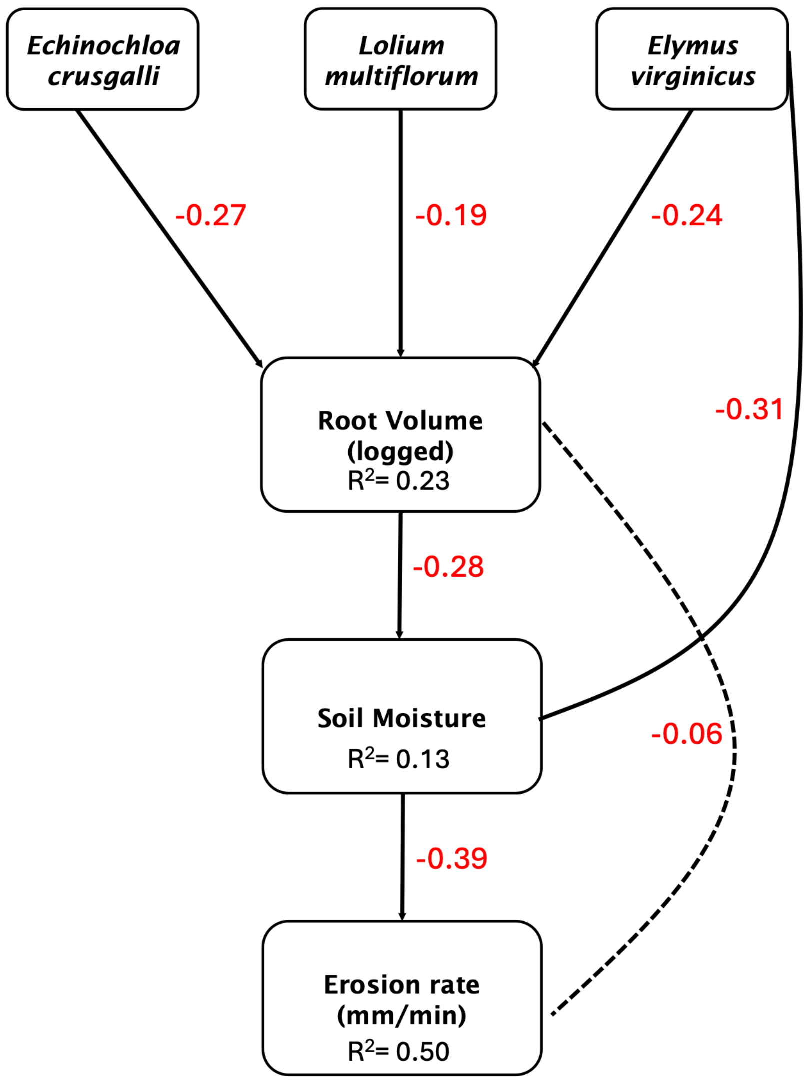


**Figure S2:** Visual for piecewise structural equation model (pSEM) for root volume.

**Table S5a:** Linear mixed effects model (LMER) of species presence effects on root volume for the first component model for the piecewise structural equation model (pSEM).

| Predictor | Response | Estimate | Std. Error | DF | t value | p value |
|---|---|---|---|---|---|---|
| *Echinochloa crus-galli* | Root volume ($mm^3$), natural-log transformed | -0.4035 | 0.1499 | 77.5 | -2.693 | **0.009** |
| *Lolium multiflorum* | Root volume ($mm^3$), natural-log transformed | -0.2906 | 0.1510 | 77.3 | -1.925 | 0.058 |
| *Elymus virginicus* | Root volume ($mm^3$), natural-log transformed | -0.3672 | 0.1489 | 74.3 | -2.467 | **0.016** |

**Table S5b:** Linear model (LM) of root volume and *Elymus virginicus* effects on soil moisture for the second component model for the piecewise structural equation model (pSEM).

| Predictor | Response | Estimate | Std. Error | DF | t value | p value |
|---|---|---|---|---|---|---|
| Root volume ($mm^3$), ln-transformed | Soil moisture content | -0.0151 | 0.0058 | 79 | -2.600 | **0.011** |
| *Elymus virginicus* | Soil moisture content | -0.0248 | 0.0086 | 79 | -2.897 | **0.005** |

**Table S5c:** Linear mixed effects model (LMER) of moisture content and root volume effects on erosion rate for the third component model for the piecewise structural equation model (pSEM).

| Predictor | Response | Estimate | Std. Error | DF | t value | p value |
|---|---|---|---|---|---|---|
| Soil moisture content | Erosion rate (mm/min) | -11.20 | 2.43 | 75.2 | -4.604 | **< 0.001** |
| Root volume ($mm^3$), ln-transformed | Erosion rate (mm/min) | -0.1011 | 0.1322 | 75.7 | -0.765 | 0.447 |

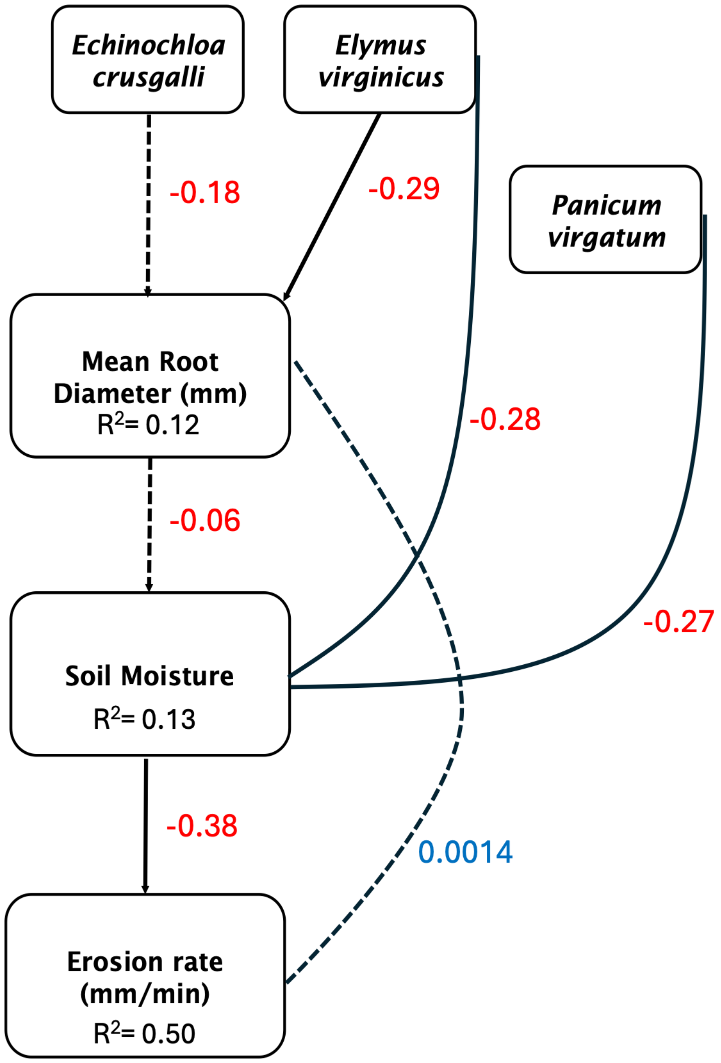


**Figure S3:** Visual for piecewise structural equation model (pSEM) for mean root diameter.

**Table S6a:** Linear model (LM) of species presence effects on mean root diameter for the first component model for the piecewise structural equation model (pSEM).

| Predictor | Response | Estimate | Std. Error | DF | t value | p value |
|---|---|---|---|---|---|---|
| *Echinochloa crus-galli* | Mean root diameter (mm) | -0.0485 | 0.0275 | 79 | -1.760 | 0.082 |
| *Elymus virginicus* | Mean root diameter (mm) | -0.0778 | 0.0279 | 79 | -2.794 | **0.007** |

**Table S6b:** Linear model (LM) of mean root diameter, *Panicum virgatum*, and *Elymus virginicus* effects on soil moisture for the second component model for the piecewise structural equation model (pSEM).

| Predictor | Response | Estimate | Std. Error | DF | t value | p value |
|---|---|---|---|---|---|---|
| Mean root diameter (mm) | Soil moisture content | -0.0180 | 0.0333 | 78 | -0.541 | 0.590 |
| *Panicum virgatum* | Soil moisture content | -0.0216 | 0.0084 | 78 | -2.568 | **0.012** |
| *Elymus virginicus* | Soil moisture content | -0.0222 | 0.0088 | 78 | -2.518 | **0.014** |

**Table S6c:** Linear mixed effects model (LMER) of moisture content and mean root diameter effects on erosion rate for the third component model for the piecewise structural equation model (pSEM).

| Predictor | Response | Estimate | Std. Error | DF | t value | p value |
|---|---|---|---|---|---|---|
| Soil moisture content | Erosion rate (mm/min) | -10.84 | 2.40 | 75.3 | -4.521 | **< 0.001** |
| Mean root diameter (mm) | Erosion rate (mm/min) | 0.0119 | 0.7161 | 75.3 | 0.017 | 0.987 |
